# Locus-specific transposable element expression drives human hematopoietic stem cell disease pathophysiology

**DOI:** 10.64898/2026.07.31.742092

**Authors:** Angelica Varesi, Sontago Dong, Chiara Gaddoni, Kerstin B. Kaufmann, Tevis Wong, Ivan Merelli, Andy G.X. Zeng, Jessica McLeod, Liqing Jin, Isabella di Biasio, Igor Novitzky-Basso, Jonas Mattson, Stephanie Z. Xie, Bowen Li, Samuele Ferrari, John E. Dick

**Affiliations:** Department of Molecular Genetics, University of Toronto, Toronto, ON, Canada M5S 1A8; Princess Margaret Cancer Centre, University Health Network, Toronto, ON, Canada M5G 1L7; Leslie Dan Faculty of Pharmacy, University of Toronto, Toronto, Ontario, Canada; San Raffaele Telethon Institute for Gene Therapy, IRCCS San Raffaele Scientific Institute, Milan, Italy; National Research Council, Institute for Biomedical Technologies, Segrate, Italy; Department of Medical Biophysics, University of Toronto, Toronto, ON, Canada M5G 1L7; Department of Medicine, University of Toronto, Toronto, ON, Canada M5S 3H2; Institute of Biomedical Engineering, University of Toronto, Toronto, Ontario, Canada; Department of Chemistry, University of Toronto, Toronto, Ontario, Canada; Vita-Salute San Raffaele University, Milan, Italy

## Abstract

VEXAS syndrome (vacuoles, E1 enzyme, X-linked, autoinflammatory, somatic) is a severe, inflammatory syndrome resulting from mutated *UBA1* leading to hematopoietic stem cells (HSC) expansion. Although *UBA1*-mutant HSC show complex phenotypes including proteostasis defects, sustained inflammation and clonal expansion of myeloid biased progeny, the pathogenic mechanisms at the HSC level are unknown from these gene-centric studies alone. By focussing on the non-coding genome and using advanced functional genetic methods, we found that VEXAS HSC, compared to controls, had altered expression of individual transposable elements (TE) and are key regulators of VEXAS pathogenesis. Locus-specific TE quantification identified two L1 elements, L1-10 and L1-15, active in both normal and VEXAS HSC that drive myeloid commitment by co-opting SPI1 and IRF1 transcription factors (TF) via networks common to other myeloid-biased conditions. Lipid nanoparticle (LNP)-mediated CRISPRi of L1-10 and L1-15 in *UBA1*-mutant HSC also caused reversion of VEXAS-associated functional phenotypes *in vitro* and *in vivo*. Functionally, pharmacologic inhibition of UBA1 with TAK-243 led to L1-10 and L1-15 RNA accumulation, while enhancement of UBA1 activity with Auranofin reversed this effect. Our study provides direct evidence that VEXAS-specific TE govern HSC clonal dominance, thereby uncovering a regulatory axis underlying HSC biology and disease mechanisms, opening a therapeutic strategy directed towards the repetitive genome.

## Main

Blood production is sustained lifelong through a unique balance where hematopoietic stem cells (HSC) maintain the HSC pool through self-renewal while still generating progressively restricted progenitors, that in turn produce a massive daily mature blood cell output ^1^. However during aging, a variety of cell-intrinsic and cell-extrinsic factors such as inflammation and somatic mutation acquisition alter this balance and act to progressively reshape the hematopoietic stem and progenitor cell (HSPC) compartment ^2,3^. This confers fitness advantage to individual HSC clones and often shifts hematopoiesis towards an oligoclonal and myeloid-biased state known as clonal hematopoiesis (CH), thereby reducing HSC pool diversity ^4–7^. Although CH is associated with increased risk of hematological malignancies and other age-related diseases, the mechanisms driving clonal expansion remain incompletely understood ^6,8–12^. Studies of clonal HSC from pre-malignant and genetic hematological disorders have provided important insights into clonal expansion and lineage skewing, yet single-cell studies often reveal surprisingly few differences between CH and healthy cells at the level of the most primitive long term (LT)-HSC ^2,13^. These observations suggest that gene-centric transcriptional changes alone may not capture the regulatory architecture underlying HSC state and clonal fitness, highlighting the importance of additional regulatory layers potentially lying in the epigenome. Transposable elements (TE) are a class of repetitive elements that comprise a large proportion of the non-coding genome. Although the expression of most TE is epigenetically silenced in somatic cells, their repression erodes during aging, cellular stress, inflammation, and cell transformation, resulting in activation of viral mimicry pathways, innate immune signaling, and senescence-associated transcriptional programs ^14–19^. Thus, TE are increasingly recognized as a non-coding-centric complementary regulatory layer of cell fate determination potentially linking aging, inflammation, stem-cell state, and aberrant hematopoietic differentiation. Beyond their transcriptional activity, TE contain cis-regulatory elements and changes in chromatin accessibility can provide binding sites for transcription factors (TF), including pioneer factors, and may contribute to higher-order genome organization through enhancer activity and chromatin looping ^20–23^. Consistent with a functional role for TE in malignant stem-cell states, we recently demonstrated that a chromatin-accessibility signature comprising 121 differential TE families (LSCTE121) stratifies patients with acute myeloid leukaemia (AML) according to prognosis; normal HSPC vs mature cell populations were also profiled ^24^. Critically, CRISPR interference (CRISPRi) targeting a single top-ranked TE family of both AML and normal datasets, LTR12C, reduced phenotypic leukaemia stem cells (LSC). These observations support a model in which TE actively shape LSC identity and cell-state transitions rather than simply reflecting global epigenetic dysregulation. However, whether TE transcriptional activity contributes to the emergence and persistence of clonally dominant non-malignant HSC remains unknown. Moreover, most studies have examined TE at the family level, despite the fact that individual families can comprise hundreds to thousands of distinct genomic loci with potentially divergent regulatory activities. The locus-specific contribution of individual TE to the biology of highly purified human HSC, particularly in a human disease that drives clonal expansion, remains unexplored.

VEXAS (vacuoles, E1 enzyme, X-linked, autoinflammatory, somatic) syndrome, a severe adult-onset inflammatory disease caused by somatic mutations in *UBA1,* is an ideal CH setting to address this question. *UBA1* encodes the major E1 enzyme that initiates ubiquitin activation and is central to protein homeostasis ^25^. *UBA1*-mutant HSPC acquire inflammatory and senescence-like programmes and show enhanced myeloid commitment, phenotypes that can be recapitulated by introducing *UBA1* mutations into human HSPC by base editing ^13,26,27^. Yet, despite profound biological consequences of *UBA1* mutations, transcriptional differences between VEXAS and healthy cells remain comparatively modest at the level of the most primitive HSC, suggesting that mutation-driven clonal dominance may involve regulatory layers that are not captured by conventional gene-centric analyses ^13,26,27^). Here, we profiled TE expression at single-cell and locus-level resolution across scRNA-seq datasets of CD34+ HSPC from healthy controls across a range of ages, from inflammation challenge models and individuals with VEXAS syndrome or pre-leukemic myelodysplastic syndrome (MDS) ^3,28^. Our findings reveal a disease-associated, locus-specific TE landscape in human HSC and identify TE activity as a previously underappreciated regulatory layer associated with VEXAS pathophysiology and more broadly with HSC state and clonal dominance.

## Results

### HSPC in patients with VEXAS syndrome show altered, disease-specific TE expression

Although there are profound alterations in the transcriptional profile of HSPC in VEXAS compared to age and sex-matched healthy donors (HD) ^13^, within the most primitive HSC compartment such differences were modest. As we recently reported on a critical role for TE in governing HSPC versus mature as well as AML LSC ^24^, we investigated whether TE might be involved in VEXAS pathobiology at the HSC level. TE were first analyzed from publicly accessible single-cell (sc)RNA-seq datasets ^13,26,28^ using the machine learning algorithm Single cell Transposable Element Locus Level Analysis of scRNA Sequencing (Stellarscope) ^29^ to compare VEXAS HSPC to age and sex-matched HD (Fig.1a). Stellarscope undertakes locus-specific TE quantification of those TE with polyA that are captured in 10x libraries (typically L1 and HERV families) and uses an expectation maximization algorithm that was previously validated in the context of human peripheral blood mononuclear cells. Multimapping reads were assigned to the most probable location and the resulting TE count matrix is corrected for overlap with exonic sequences, thus avoiding confounding artifacts from canonical gene expression ^29^. Amongst all TE elements assessed with Stellarscope, only a limited number of loci belonging to the L1 and HERVs families distinguished HSPC subtypes found in VEXAS samples as compared to HD (Extended Data Fig.1a). Differential TE expression analysis revealed a limited and variable number of TE loci significantly up- or down-regulated (between 63 and 2, respectively) across VEXAS hematopoietic HSPC cell types as previously defined in ^13^ (Fig.1b). Most of the loci differentially expressed in VEXAS HSPC (∼80%) belong to L1 (annotated as L1FLnI or L1ORF2) that can no longer undergo reverse transcription ^29^. Less than ∼20% differentially expressed individual loci are HERVs or L1 elements annotated as L1FLI with retrotransposition potential (Fig.1b and Extended Data Fig.1b). Intriguingly, the most primitive HSPC subsets (HSC, MPP/CMP and Myelo/Lympho/CMP) had the highest number of differentially expressed individual loci considering either all of them (L1FLI and HERVs) or L1FLI-only (Fig. 1b and Supplementary Table 1). When focusing on the 21 TE differentially expressed in the HSC cluster, we observed that fold change in VEXAS versus HD varied across the hematopoietic hierarchy, with some TE being upregulated in primitive VEXAS populations and downregulated in differentiated cells, and vice versa (Extended Data Fig.1c). Two active L1 elements residing in distal intergenic regions – L1FLI-15q21.3 and L1FLI-10q25.1b (hereafter referred to as L1-10 and L1-15) – were the most upregulated in VEXAS HSC and MPP/CMP compared to HD (Fig.1c). Their fold change in VEXAS versus HD was positive in the most primitive HSPC compartment, but progressively changed into negative values upon lineage commitment (Extended Data Fig.1c and Extended Data Fig.1d-g). To evaluate the extent to which the TE differentially expressed in VEXAS versus HD HSC were also stem cell specific, we defined a VEXAS TE Primitive Score by calculating their pseudotime-weighted average expression difference in VEXAS HSC versus committed populations. Accordingly, the two active L1 elements, together with the inactive L1FLnI-13q31.2d, remained the most upregulated and specific in VEXAS HSC (Fig.1d). Overall, these data reveal extensive remodeling of the TE landscape in VEXAS HSPC, characterized by inverse expression patterns between primitive and committed compartments.

**Fig. 1.**
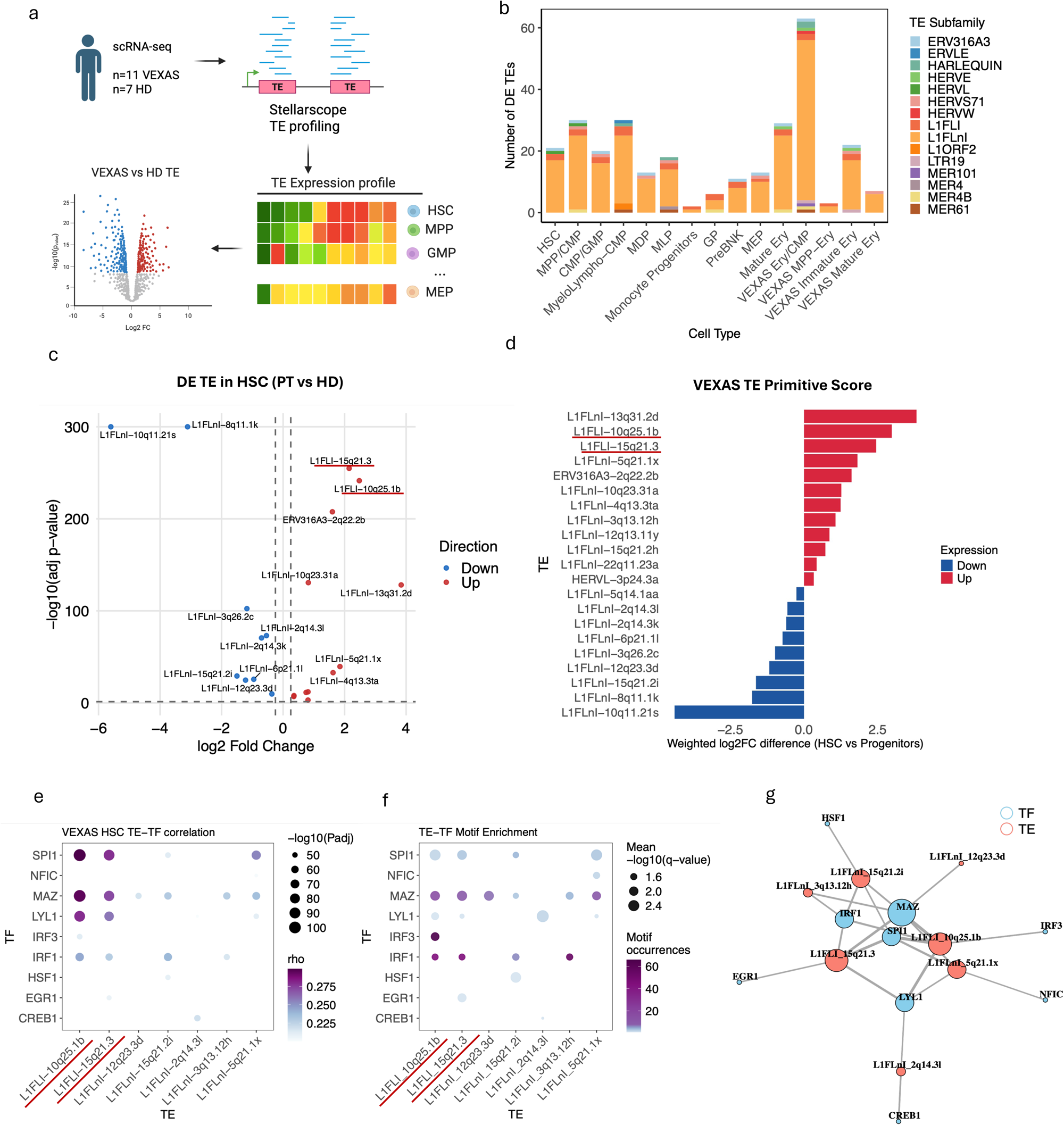
A VEXAS-specific TE expression landscape in HSPC. a) Scheme of the scRNA-seq cohort and bioinformatic analysis. b) Number of differentially expressed TE loci in VEXAS patients versus healthy donors (HD) across HSPC populations as defined in ^13^. Colors denote TE family. c) Volcano plot showing upregulated (red) and downregulated (blue) TE loci in VEXAS HSC versus HD. The two active L1 elements mostly enriched in VEXAS HSC versus HD are underlined in red. d) Bar plot showing the pseudotime-weighted log2 fold-change difference between HSC and all other HSPC populations for each of the 21 TE loci differentially expressed in VEXAS HSC shown in (c) e) Dot plot showing Spearman correlations between TE expression and TF expression in VEXAS HSC. Only significant TE–TF pairs are displayed. f) Dot plot showing the number of predicted TF motif binding sites within TE loci for the significant TE–TF pairs shown in (e). g) TE–TF correlation network obtained from the significant Spearman correlations shown in (e). Nodes represent TE loci and TF, with node size proportional to the number of significant TE–TF correlations (node degree). Edges connect significantly correlated TE–TF pairs, and edge width is proportional to the significance of the correlation (−log10 adjusted *P* value).

### L1-10 and L1-15 reside in a core VEXAS-associated TE-TF regulatory network

TE sequences often harbor TF binding sites and represent a source of cis-regulatory potential ^30^. To investigate the cis-regulatory relevance of the TE in VEXAS, and to understand whether TE expression contributes to HSC clonal dominance, we focused on the 21 TE differentially expressed in VEXAS HSC vs HD (VEXAS TE Primitive Score) and correlated their expression with the expression of TF in patient HSC. After filtering (Methods), nine TF and seven TE were highly intercorrelated (Fig.1e). Of note, L1-10 and L1-15 showed high correlation with SPI1 (also known as PU.1), MAZ, LYL1, and IRF1 (Fig. 1e). Additionally, IRF1, IRF3 and MAZ also showed a considerable number of motif occurrences (>20) in the sequence of the correlated TE (Fig. 1f). The resulting TE-TF network showed a tight interconnection between L1-10, L1-15, L1FLnI-5q21.1x, L1FLnI-15q21.2i, SPI1, LYL1, MAZ and IRF1, with L1-10 and L1-15 displaying the highest level of significance and the highest number of connections (Fig. 1g). Of note, the components of this network are well-known regulators in hematopoiesis and stemness: SPI1 encodes for PU.1, the master factor governing myeloid differentiation; IRF1 is a critical TF that acts as a bridge between inflammatory signaling and stem cell maintenance, and MAZ and LYL1 contribute to the regulation of genes required for HSC activation and stem cell maintenance. Given that enhanced interferon signaling and myeloid skewing are hallmarks of VEXAS HSPC, our findings raise the possibility that TE dysregulation contributes to disease pathogenesis through co-opting and reinforcing defined TE–TF regulatory networks. To carry out a more detailed investigation of this possibility, we performed an unsupervised analysis of clusters of highly correlated genes in VEXAS and HD HSC. Weighted Gene Co-expression Network Analysis (hdWGCNA) was used to identify clusters of highly connected regulators (“hub genes”) that control entire pathways. hdWGCNA resulted in 5 gene modules characterizing VEXAS HSC (PT-M1 to PT-M5) and an additional 5 modules for healthy HSC (HD-M1 to HD-M5). Except PT-M3, which showed partial overlapping with HD-M1, there was minimal similarity between genes and pathways enriched among all the modules (Methods; Extended Data Fig.2a,b). Gene ontology enrichment analysis identified distinct biological pathways across these HD and PT gene modules (Extended Data Fig.2c,d). Among the most enriched processes in VEXAS HSC modules were mitochondria metabolism, cell replication, response to DNA damage, antiviral response, protein folding, and antioxidant responses, in line with the increased accumulation of unfolded proteins and cellular stresses in this disease (Supplementary Table2 and Supplementary Table3). These VEXAS specific modules are also shared with other inflammatory and/or hematological conditions including those from (i) Zeng et al. ^3^, which describes two distinct HSC subsets, i.e. homeostatic HSC and HSC with molecular memory of prior inflammation (HSC-iM); (ii) Ainciburu et al. ^28^, which compares HSC from elderly and young HD, and from MDS patients and age-matched HD, (iii) Cheong et al ^31^, which investigates HSC from HD and following COVID-19 infection, and (iv) Jakobsen et al ^2^, which compares HSC from TET2 and DNMT3A mutant clonal hematopoiesis donors with healthy HSC (Extended Data Fig.2e-g). Transcriptional overlap using hdWGCNA module preservation analysis ^32^ all VEXAS PT modules were moderately to highly conserved, with PT-M1, PT-M2 and PT-M3 being the most maintained throughout datasets and conditions irrespective from sex (Extended Data Fig.2e-g). These data indicate VEXAS HSC are characterized by the co-existence of 5 transcriptional programs, which are largely unique to VEXAS compared to healthy donors but substantially overlap with gene expression features of similar pro-inflammatory and age-related diseases.

To determine whether VEXAS primitive TE and their associated TF are integrated into coordinated transcriptional programs, we utilized a strategy described by Colombo et al. ^33^ where the expression of TF and TE were correlated with hdWGCNA gene programs. There was generally strong correlation across VEXAS TE and PT M2-M5, but not HD M2-M5, and this was particularly marked for L1-10 and L1-15 (Extended Data Fig.3a). Similarly, we found a significant correlation between our TF of interest and PT modules, but not with HD modules (Extended Data Fig.3b). In both cases, HD-M1 was an exception, with significant correlation occurring but limited to 5 out of 21 VEXAS TE and to the TF SPI1, LYL1 and MAZ (3 out of 9; Extended Data Fig.3a,b). This is consistent with the partial overlap of HD-M1 and PT-M3. In contrast to the shared transcriptional programs between VEXAS and similar disease conditions, there was minimal to no correlation between PT M1-M5 and the TF predicted to be co-opted by VEXAS primitive TE (Extended Data Fig.3c-f). These data suggest that despite sharing transcriptional programs with several pathophysiological hematological conditions HSC from VEXAS patients have a unique TE-TF network.

### A unique TE signature distinguishes VEXAS HSC from those of related hematological states

Next, we assessed whether the transcriptional similarities found between HSC from VEXAS patients and HSC from other broadly related pathophysiological conditions also extend to the non-coding genome of these related HSC cell types. Given the limitations imposed by Stellarscope’s input requirements for raw sequencing files, we restricted our analysis to scRNAseq from Zeng et al and Ainciburu et al., which enabled us to profile TE expression in HSC with memory of past inflammatory exposure, MDS HSC, and aged HSC. There were 62 single TE loci differentially expressed in HSC harboring memory of past inflammatory exposure (HSC-iM) compared to homeostatic HSC (HSC-I) (Extended Data Fig.3g). 60 out of 62 belonged to the L1FLnI TE subfamily, and only 2 to HERVs (Extended Data Fig.3g and Supplementary Table4). Of note, no active L1 were found to be differentially expressed. 2 TE residing in close proximity on the q arm of chr7 - namely L1FnI-7q33u and L1FLnI-7q34a - were distinctively higher expressed in HSC-iM versus HSC-I (Extended Data Fig.3g). MDS HSC versus old HSC had 14 differentially expressed TE loci, half of which were upregulated and the rest downregulated (Extended Data Fig.3h). All of them except one belonged to the non-active L1FLnI TE subfamily (Extended Data Fig.3h and Supplementary Table5). L1FLnI-5p15.2o and L1FLnI-5q15k were the 2 most differentially upregulated, while L1FLnI-10q11.21f and L1FLnI-15q21.2i were the top 2 most downregulated (Extended Data Fig.3h). While L1FLnI-5p15.2o is distal intergenic and L1FLnI-5q15k resides in the intronic region of *ARB2A* gene, L1FLnI-10q11.21f overlaps with a *CXCL12* exon, a critical chemokine frequently dysregulated in MDS HSPC, and L1FLnI-15q21.2i is in the promoter region of *SSBP2*, which exerts tumor suppression roles in leukemia transformation ^34^. (Supplementary Table6). Finally, only 12 differentially expressed TE were found when comparing aged versus young HSC (Extended Data Fig.3i). All of them except one were inactive L1 elements (Extended Data Fig.3i). Importantly, the only active L1 was L1-15, one of the two TE loci most upregulated in VEXAS HSC versus HD HSC (Extended Data Fig.3i and Fig.1c).

We then sought to quantify how many TE loci from the VEXAS TE primitive score were shared with MDS, aged or inflamed HSC. Strikingly, only a minority of the differentially expressed TE in VEXAS HSC versus HD were also significantly different in MDS vs HD HSC (9), Aged vs Young HSC (8), or HSC-iM vs HSC-I (7) (Fig. 2a-c). Despite reaching significance, differential expression of the top-ranking VEXAS HSC TE was either negligible or negatively enriched across all the above comparisons (Fig.2d). To quantify the extent to which the top VEXAS TE are also unique to VEXAS, we calculated a VEXAS TE Unique Score by computing the log2FC difference between VEXAS HSC and each of the following comparisons, as before: MDS vs HD HSC, Aged vs Young HSC, or HSC-iM vs HSC-I. This analysis confirmed that the top most differentially expressed VEXAS TE, including L1-10 and L1-15, were specific to VEXAS (Fig. 2e-g). Overall, these findings indicate that VEXAS HSC possess a unique TE landscape compared to HSC from related hematologic conditions, despite some comparability of their transcriptional and health states.

**Fig. 2.**
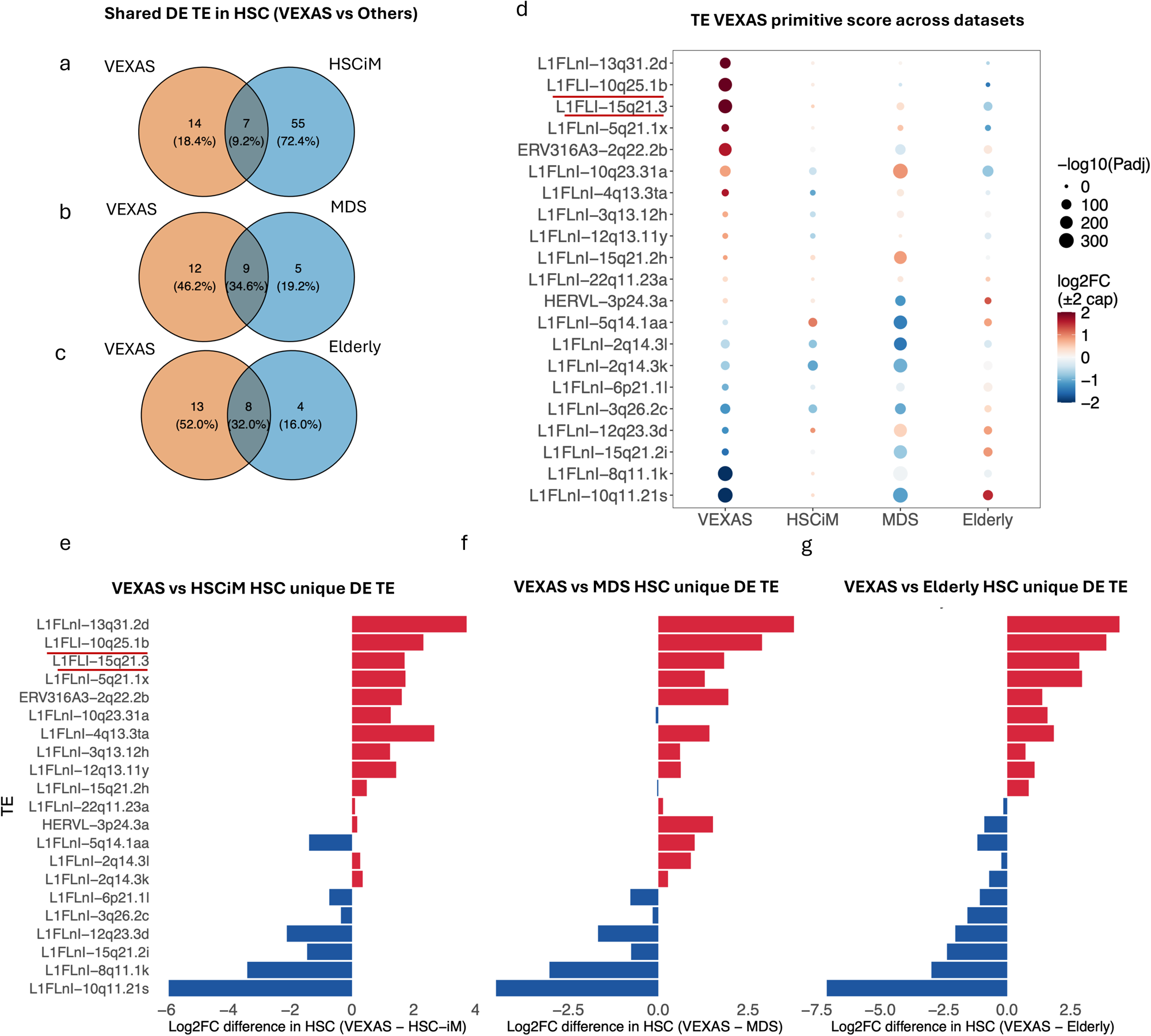
Distinct TE landscapes define VEXAS HSPC and related hematological conditions. a-c) Venn diagrams showing the overlap between TE loci differentially expressed in VEXAS HSC relative to HD and in (a) HSC-iM relative to HSC-I, (b) MDS HSC relative to healthy HSC, and (c) elderly HSC relative to young HSC. d) Dot plot showing differential expression of the 21 TE loci identified in Fig. 1c in VEXAS HSC relative to HD; HSC-iM relative to HSC-I; MDS HSC relative to healthy HSC, and elderly HSC relative to young HSC. e-g) Bar plots showing the log2 fold-change difference of the 21 TE loci identified in Fig. 1c between VEXAS HSC relative to HD and (e) HSC-iM relative to HSC-I, (f) MDS HSC relative to healthy HSC, and (g) elderly HSC relative to young HSC.

### L1-10 and L1-15 upregulation induces myeloid bias and VEXAS-like features in wild-type HSPC

Our computational data proposes that L1-10 and L1-15 might influence HSC biology and be linked to key VEXAS-like biological properties including myeloid bias, so we undertook functional studies to obtain direct evidence for this hypothesis. Using an optimized CRISPR activation (CRISPRa) gene editing approach, L1-10 and L1-15 were individually transactivated in mPB CD34+ HSPC from healthy male donors by mRNA delivery of a dCas9-VPR and pairs of gRNA (Fig.3a and Supplementary Table7). To ensure locus specific transactivation and minimize off targets, the gRNA were designed to the most proximal unique region upstream of the target TE, thus minimizing the challenge of targeting highly similar repetitive regions (Fig.3a). Non-targeting neutral gRNAagainst the safe harbor *AAVS1* locus was used as a control in all our experiments ^35^. The efficiency and specificity of transactivation was evaluated using specific primers amplifying each target TE transcript and quantifying their abundance by digital droplet PCR (ddPCR) 48h and 96h after editing (Fig.3a). Electroporation of gRNA pairs targeting L1-10 or L1-15 together with dCas9 VPR in mPB CD34+ HSPC led to a specific transactivation of L1-10 or L1-15 48h and 96h compared to *AAVS1* control (Fig 3 b,c and Extended Data Fig.4a,b). As predicted from our *in silico* analysis, TE transactivation in HSPC at 48h led to the overexpression of the SPI1 and IRF1 TF (Extended Data Fig.4c,d). This was similar at 96h, indicating that L1-10 and L1-15 transactivation may be sufficient to feed-forward activate the expression of TF whose binding site is embedded within them (Fig.3d,e). SPI1 and IRF1 upregulation as a direct consequence of L1-10 and L1-15 transactivation was intriguing, as this mirrors the transcriptional landscape of VEXAS HSC characterized by myeloid skewing and enhanced interferon type-I transcriptional response ^13^.

**Fig. 3.**
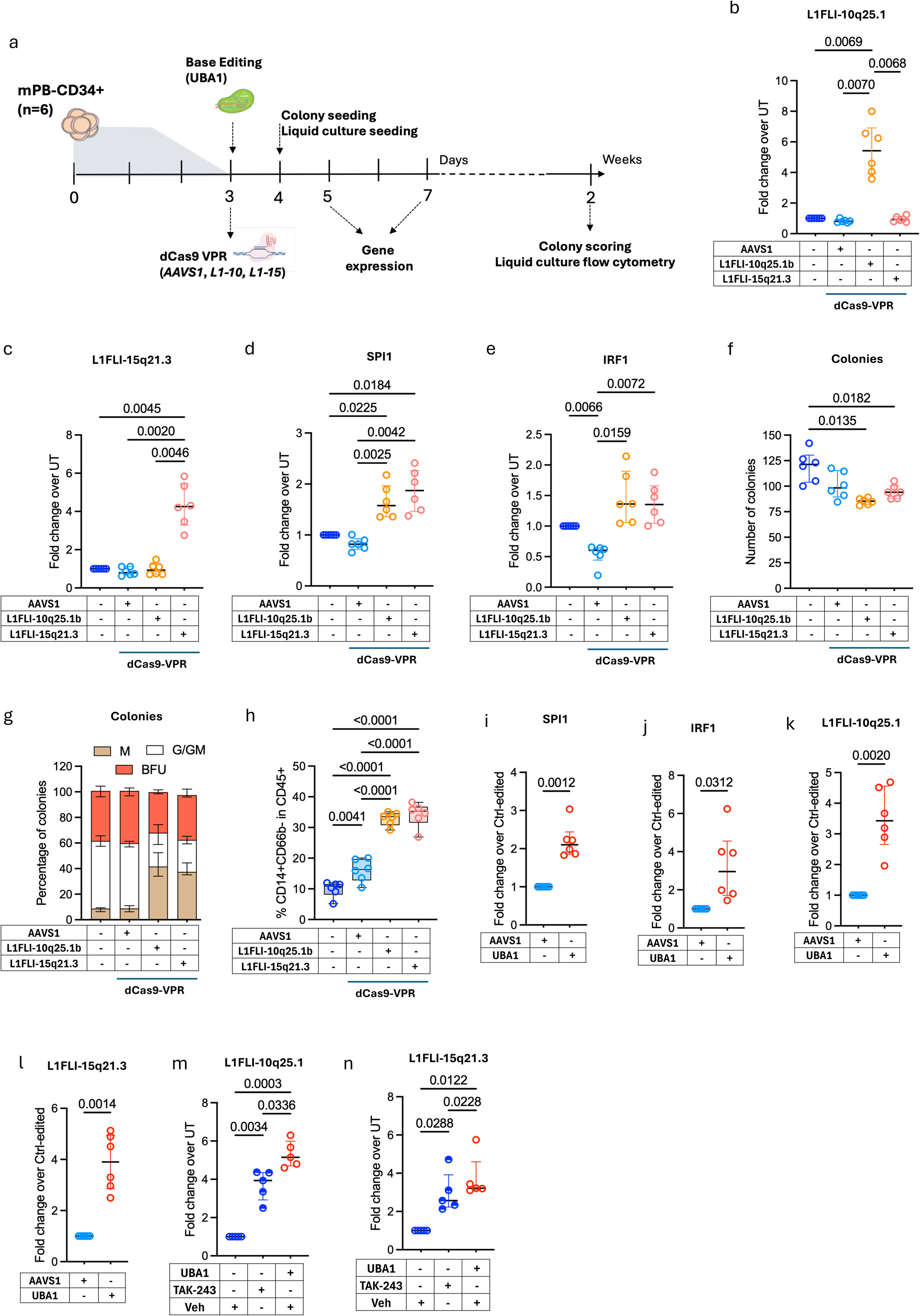
L1-10 and L1-15 transactivation induce VEXAS-like myeloid skewing in adult HSPC. a) Scheme of the CRISPRa editing strategy. b-e) Fold change in expression of (b) L1-10, (c) L1-15, (d) SPI1, and (e) IRF1 of HSPC edited as indicated relative to untreated cells (UT) 96 h after editing, measured by ddPCR (n=6). f) Number of colonies obtained on seeding HSPC edited as indicated (n=6). g) Proportion of myeloid (M), granulocytic/granulocytic-myeloid (G/GM) and erythroid (BFU) colonies from the condition in (f) (n=6). h) Proportion of CD14+CD66b-within CD45+ cells measured by flow cytometry of HSPC edited as indicated after two weeks of differentiation culture (n=6). i-l) Fold change in expression of (i) SPI1, (j) IRF1, (k) L1-10, and (l) L1-15 of *UBA1* mutant HSPC relative to control edited cells 96 h after editing, measured by ddPCR (n=6). m,n) Fold change in expression of (m) L1-10, and (n) L1-15 of *UBA1* mutant HSPC or HSPC treated with TAK-243 (10nM) relative to vehicle treated cells 96 h after treatment/editing, measured by ddPCR (n=5).

To determine whether L1-10 and L1-15 upregulation led to phenotypic consequences, we focused on methylcellulose clonogenic assays. L1-10 and L1-15 upregulation did not impact total clonogenicity as compared to the *AAVS1* control (Fig.3f). However, transactivation of either, in healthy HSPC, was sufficient to significantly increase the proportion of myeloid colonies (∼45% of CFU) compared to neutral controls (∼7% of CFU) (Fig.3g). These data were independently corroborated by flow cytometric analysis of cells cultured for 2 weeks in differentiation conditions ^36^. L1-10 and L1-15 transactivation markedly increased the proportion of CD14+ monocytes (∼35% compared to ∼16% in *AAVS1* controls) without impacting the fraction of CD34+ stem cells (Fig.3h and Extended Data Fig.4e). Given the similarity with the phenotype observed in VEXAS patients and in vitro models, these data raised the possibility that activation of these specific L1 elements might occur as consequence of the *UBA1* mutation in HSC and this results in reshaping of the VEXAS stem cell state and myeloid bias through the induction of key transcriptional regulators. To test this hypothesis, we modelled VEXAS syndrome by inserting the most common mutation (p.Met41Thr) in healthy male mPB CD34+ HSPC by mRNA delivery of the ABE8.20-m base editor together with a gRNA targeting Met41, as previously reported ^13^. The editing efficiency ranged from 80-90% across all donors (Extended Data Fig.4f,g). In line with increased myeloid skewing and inflammatory transcriptional signatures reported in patient samples, *UBA1* mutant CD34+ cells showed higher expression of SPI1 and IRF1 at both 48h and 96h after editing (Fig.3i,j and Extended Data Fig.4h,i). As previously reported, methylcellulose assays of *UBA1* mutant HSPC showed a significantly reduced clonogenic output and an almost exclusive differentiation in myeloid colonies ^13^ (Extended Data Fig.4j,k). Finally, we found that L1-10 and L1-15 expression was elevated by 3-4 fold at the 96h timepoint only in the *UBA1* mutant HSPC group providing strong evidence that locus-specific dysregulated TE expression in VEXAS HSC arises as a direct consequence of *UBA1* mutation and loss of function (Fig.3k,l).

To gain independent evidence that the increase in L1-10 and L1-15 expression was due to the general consequence of UBA1 loss of function rather than some feature of UBA1 base editing, we treated wild type HSPC with TAK-243, a targeted UBA1 inhibitor. TAK-243 treatment for 96 hours was sufficient to significantly induce L1-10 and L1-15 transcription, albeit at a lower level compared to *UBA1* mutant HSPC (Fig.3m,n). Colony assessment at 2 weeks showed a consistent reduction of the number of colonies in WT HSPC treated with TAK-243 to levels very close to *UBA1* mutant HSPC (Extended Data Fig.4l). In both cases, the resulting colonies were predominantly myeloid (∼60% in *UBA1* mutant HSPC and ∼50% in TAK-243 treated healthy HSPC), indicating that loss of UBA1 activity is sufficient to induce myeloid bias and cause L1-10 and L1-15 upregulation (Extended Data Fig.4m). Together, these findings, using two independent approaches, identify L1-10 and L1-15 as prominent features of the VEXAS HSC state, show that their activation is sufficient to recapitulate key transcriptional and functional characteristics of VEXAS hematopoiesis, and demonstrate that their upregulation is consistently associated with genetic or pharmacological inhibition of UBA1 activity.

### L1-10 and L1-15 silencing in *UBA1* mutant HSPC partially rescues VEXAS phenotypes

To further link L1-10 and L1-15 with the HSPC phenotype exerted by *UBA1* mutation in VEXAS, we undertook the orthogonal approach of silencing L1-10 and L1-15 in *UBA1* mutant HSPC to determine if the phenotype can be reverted. The second goal of such a study is to validate the possibility that L1-10 and L1-15 silencing might represent a therapeutically actionable target in VEXAS. As a first step to validate this approach, we first tested the impact of L1-10 and L1-15 silencing in healthy male mPB HSPC using a CRISPR inactivation strategy (CRISPRi) involving the delivery of dCas9-KRAB together with the same pairs of gRNA used for the CRISPRa experiments. L1-10 or L1-15 expression was significantly decreased (∼4-fold) 48h after CRISPRi and remained reduced to similar levels at 96h (Extended Data Fig.5a-d). Similar to the CRISPRa experiments, this approach exhibited high target specificity, as repression of each TE locus selectively affected its intended target without altering the expression of the other. Interestingly, L1-10 or L1-15 silencing did not impact the expression of SPI1 and IRF1 at either timepoints (Extended Data Fig.5e-h). Furthermore, CRISPRi of either TE did not alter total clonogenic capacity or multilineage clonogenic ability of edited HSPC (Extended Data Fig.5i,j). In line with these data, L1-10 or L1-15 silencing did not impact the proportion of CD34+ nor the fraction of CD14+ cells after 2 weeks of liquid culture in differentiation conditions (Extended Data Fig.5k,l). Overall, these findings indicate that CRISPRi of L1-10 and L1-15 does not alter the phenotype of healthy mPB HSPC *in vitro*.

To test the impact of L1-10 and L1-15 silencing in VEXAS HSPC, the *UBA1* mutation was introduced into male mPB using base editing and then CRISPRi for L1-10 and L1-15 was subsequently performed. We had to develop an optimized approach that involved LNP delivery of the L1-10 and L1-15 CRISPRi mRNA as there is high toxicity of sequential electroporation ^37^. LNP have lower toxicity and similar efficiency compared to electroporation ^38^. We first showed that there was strong silencing of L1-10 or L1-15 upon transfection with LNP containing gRNAs and dCas9-KRAB mRNA (Extended Data Fig.5m,n). As in our previous experiments, we did not notice any phenotypic difference between L1-10 and L1-15 perturbation, therefore we reasoned that delivering a mix of LNP simultaneously targeting L1-10 and L1-15 would be of therapeutic advantage. Of note, cell transfection with half dose of LNP targeting L1-10 and half targeting L1-15 yielded comparable transactivation or downregulation efficiency than individual targets alone (Extended Data Fig.5m,n). We then optimized a sequential editing protocol to introduce the *UBA1* mutation by electroporation at day 2 followed by TE silencing by LNP at day 3, targeting L1-10 and L1-15 simultaneously (Extended Data Fig.6a). The increased expression of L1-10 and L1-15 at 96h post editing in *UBA1* mutant HSPC receiving *AAVS1*-targeting CRISPRi LNP was confirmed (Fig.4a,b). By contrast, L1-10 and L1-15 expression was reduced by 88% in *UBA1* mutant HSPC receiving the L1-10 and L1-15 targeting CRISPRi LNP as compared to unedited controls (Fig.4a,b). SPI1 and IRF1 followed a similar pattern of expression: while both TF were upregulated in *UBA1* mutant HSPC receiving control LNP (∼ranging between 2 and 4 fold), their expression was reduced to levels comparable with untreated cells (1.5 fold for SPI1 and 1 fold for IRF1 at 96h post editing) (Fig.4c,d and Fis.S6b,c) We next asked whether this reduced SPI1 and IRF1 TF expression, as a consequence of re-establishing basal expression of L1-10, L1-1,5, would functionally rescue the phenotypic consequences of *UBA1* mutation *in vitro*. Strikingly, CRISPRi of L1-10 and L1-15 in *UBA1* mutant HSPC doubled their clonogenic potential (∼50 colonies in *UBA1* mutant HSPC treated with L1-10 and L1-15 LNP versus ∼25 colonies in the control condition and ∼95 in untreated cells) (Fig.4e). Furthermore, while ∼70% of the colonies from *UBA1* mutant HSPC treated with control LNP were myeloid, *UBA1* mutant HSPC treated with L1-10 and L1-15 CRISPRi LNP were multilineage, (∼20% BFU, ∼70% G/GM and ∼10% M), and their colony output was similar to that of untreated cells (Fig.4f). Flow cytometric analysis of 2 week liquid cultures showed that L1-10 and L1-15 silencing doubled the proportion of CD34+ cells (∼from 18% to ∼32%) and drastically reduced (from ∼45% to ∼18%) the percentage of CD14+CD66b-monocytes in *UBA1* mutant HSPC (Fig. 4g,h). Overall, these data indicate that L1-10 and L1-15 repression partially rescues the VEXAS phenotype *in vitro*.

**Fig. 4.**
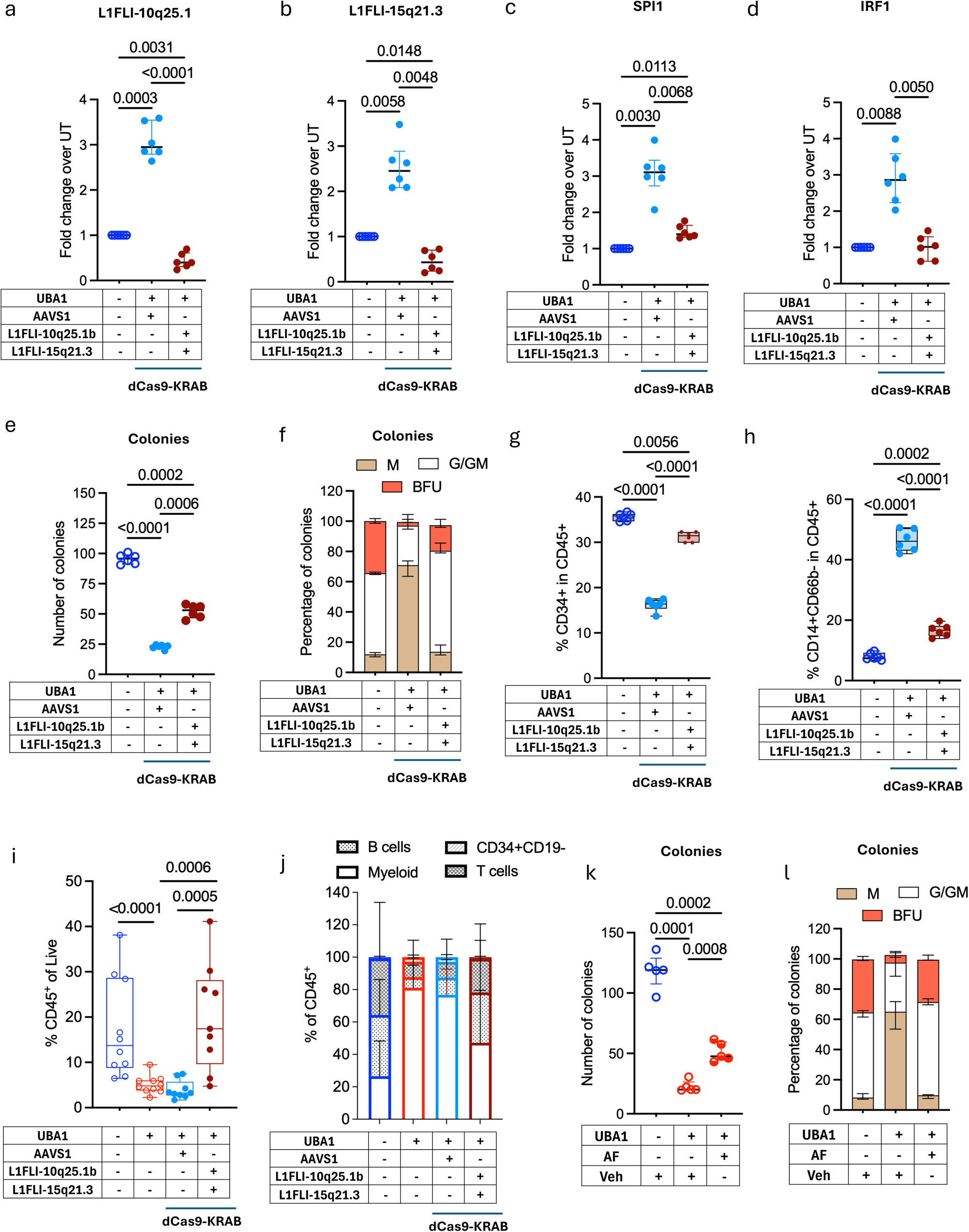
L1-10 and L1-15 repression mitigates *UBA1*-mutant HSPC phenotypes *in vitro* and *in vivo*. a-d) Fold change in expression of (a) L1-10, (b) L1-15, (c) SPI1, and (d) IRF1 of HSPC edited as indicated relative to UT 96 h after editing, measured by ddPCR (n=6). e) Number of colonies obtained on seeding HSPC edited as indicated (n=6). f) Proportion of myeloid (M), granulocytic/granulocytic-myeloid (G/GM) and erythroid (BFU) colonies from the condition in (e) (n=6). g,h) Proportion of (g) CD34+ and (h) CD14+CD66b-within CD45+ cells measured by flow cytometry of HSPC edited as indicated after two weeks of differentiation culture (n=6). i) Percentage of CD45+ human cells in the bone marrow of xenografted mice at 8w postengraftment. j) Proportion of CD33+CD19- (Myeloid), CD19+ (B cells), CD3+ (T cells) and CD34+CD19- (CD34+CD19-) cells within CD45+ human cells shown in (i). k) Number of colonies obtained on seeding *UBA1* mutant HSPC and *UBA1* mutant HSPC treated with Auranofin (AF, 100nM) relative to vehicle treated cells (n=5). l) Proportion of myeloid (M), granulocytic/granulocytic-myeloid (G/GM) and erythroid (BFU) colonies from the condition in (k) (n=5).

Finally, we turned to xenografting to directly test whether L1-10 and L1-15 silencing in *UBA1* mutant HSPC is sufficient to rescue VEXAS stemness-related phenotypes (Extended Data Fig.6d). NSG-SGM3 mice were transplanted with *UBA1* mutant HSPC with or without L1-10-and L1-15- or *AAVS1*-targeting CRISPRi LNP, along with untreated controls. As expected, *UBA1* mutant HSPC gave rise to a significantly reduced graft compared to untreated cells (∼5% versus 10-38%) (Fig. 4i and Extended Data Fig.6e). Strikingly, LNP-mediated CRISPRi of L1-10 and L1-15 - but not *AAVS1* was sufficient to significantly increase engraftment of *UBA1* mutant HSPC to levels comparable to untreated controls (Fig. 4i and Extended Data Fig.6e). Lineage analysis showed that ∼80% of the graft from *UBA1* mutant HSPC was composed of CD33+ myeloid cells, while only a minority were B, T or CD34+CD19-lineages (Fig. 4j and Extended Data Fig.6f-i). This was similar in *UBA1* mutant HSPC treated with CRISPRi LNP against *AAVS1*. As expected, xenografts from untreated HSPC showed a balanced lineage output constituted by ∼25% myeloid cells, 1% of CD34+CD19-cells, ∼34% T cells, and the remaining ∼40% of B cells (Fig. 4j and Extended Data Fig.6f-i). LNP-mediated CRISPRi of L1-10 and L1-15 shifted lineage output towards that of untreated HSPC, yielding a multilineage graft with substantially reduced myeloid predominance (∼46%) and increased lymphoid contribution (∼34% B cells and ∼20% T cells) compared to *UBA1* mutant HSPC. (Fig. 4j and Extended Data Fig.6f-i). Together, these findings show that LNP-mediated silencing of L1-10 and L1-15 restores both engraftment and multilineage hematopoiesis of *UBA1* mutant HSPC *in vivo*.

Although genetic silencing of L1-10 and L1-15 ameliorated the VEXAS phenotype in vitro and in vivo, the clinical implementation of this approach remains limited by the challenges associated with autologous hematopoietic stem cell transplantation in VEXAS patients and the absence of efficient lipid nanoparticle platforms for selective *in vivo* delivery to human HSPC. Auranofin (AF) is a clinically approved pleiotropic rheumatoid arthritis drug which has been shown to enhance UBA1 activity ^39^. Given that genetic or pharmacologically-mediated loss of UBA1 activity leads to L1-10 and L1-15 activation and mimics VEXAS phenotype, we reasoned that enhancing UBA1 activity through AF may counteract these effects, resulting in reduced L1-10 and L1-15 expression and diminished myeloid bias. Treatment of *UBA1* mutant HSPC with AF significantly reduced L1-10 and L1-15 expression at 96 post treatment compared to *UBA1* mutant HSPC in vehicle conditions (Extended Data Fig.6j,k). Additionally, AF treatment increased the total clonogenic capacity and restored the multilineage differentiation ability of *UBA1* mutant HSPC (Fig. 4k,l). Collectively, these findings demonstrate that both genetic silencing of L1-10 and L1-15 and pharmacological enhancement of UBA1 activity converge to suppress TE dysregulation and restore key functional defects of *UBA1* mutant HSPC, including impaired engraftment, myeloid skewing, and reduced multilineage potential.

## Discussion

Our study identifies locus-specific expression of distinct transposable elements as a previously unrecognized regulatory mechanism governing the human stem cell state and disease pathophysiology. By integrating single-cell transcriptomics with locus-specific TE quantification and functional perturbation, we show that two LINE1 elements are selectively activated in UBA1-mutant HSC and contribute to pathophysiological phenotypes of VEXAS syndrome, including myeloid bias, altered stem-cell fitness, and clonal expansion. These findings extend our current understanding of VEXAS pathogenesis to the non-coding genome and establish a direct mechanistic link between expressed individual transposable elements and human stem-cell disease.

By selectively modulating the most dysregulated TE loci in VEXAS HSC, we establish a genetic strategy to interrogate TE function at single-locus resolution. This represents a substantial advance over previous approaches targeting entire TE families or subfamilies, as these will simultaneously perturb thousands of highly repetitive genomic copies and obscure the contribution of individual loci ^24,40^. Our findings demonstrate that specific TE loci rather than TE subfamilies as a whole are sufficient to drive disease-associated TE activity, highlighting the potential for locus-specific TE perturbation becoming a more precise framework for dissecting the biological relevance of the non-coding genome across many human disease settings. Moreover, we also found phenotypic consequences of L1-10 and L1-15 transactivation in normal HSPC pointing to a generalized role that TE may have in HSC regulation. Our data further suggests that TE act as cis-regulatory elements, where transcription of distal intergenic TE loci promotes the expression of master regulators of lineage commitment and inflammatory programs, including SPI1 and IRF1. Given that L1-10 and L1-15 transactivation in normal HSC induces a myeloid bias without impairing *in vitro* clonogenic capacity or multilineage differentiation, the concomitant presence of *UBA1* mutation in VEXAS may be a prerequisite for a self-reinforcing regulatory circuit. In this model, L1-10 and L1-15 induce SPI1 and IRF1 expression, whereas IRF factors driven by mutated UBA1 during stress signaling in turn promote L1 transcription ^41–43^, establishing a positive feedback loop that ultimately drives the expansion of *UBA1*-mutant HSC. Whether this circuitry also involves reverse transcription or retrotransposition will require future investigation.

Our study offers an alternative therapeutic avenue for the treatment of VEXAS patients where L1-10 and L1-15 serve as potential targets. Their selective requirement in *UBA1*-mutant HSC suggests a potential therapeutic window in which co-incident normal HSC would be spared. We show that silencing these loci early after acquisition of *UBA1* mutation attenuates the VEXAS phenotype, although whether this feedback loop remains reversible at later disease stages requires investigation in patient-derived models. Our studies provide a blueprint for clinical translation either through Auranofin mediated enhancement of L1-10 and L1-15 expression or through direct LNP mediated locus specific TE targeting. Finally, the finding that locus-specific TE activity can shape stem-cell state, influence gene-regulatory network and contribute to disease should serve to draw further attention to the role that the non-coding genome plays in health and disease and the need to integrate this perspective alongside traditional gene-centric approaches.

## Methods

### Ethics statement

Human mPB samples were obtained with informed consent from Hans Messner Allogeneic Transplant Program via the Leukemia Tissue Bank at Princess Margaret Cancer (Toronto, ON) according to procedures approved by the University Health Network (UHN) Research Ethics Board (REB# 02-0763). Research using human material was performed in accordance with all relevant guidelines and regulations.

### Primary samples

Primary samples from G-CSF-treated male, unless otherwise specified, mPB donors were obtained from the Hans Messner Allogeneic Transplant Program via the Leukemia Tissue Bank at Princess Margaret Cancer. CD34+ HSPC were purified with the CD34+ MicroBead kit (Miltenyi), following manufacturer’s instructions. Cells were cryopreserved at -150 °C.

### Plasmids, gRNAs and mRNA IVT

The ABE8.20-m plasmid for mRNA in vitro transcription (IVT) was generated in ^1^.The dCas9-KRAB and dCas9-VPR plasmids for mRNA IVT were generated in ^2^. The *AAVS1* gRNA used as edited control was previously reported ^2^. The *UBA1* gRNA introducing the p.Met41Thr mutation by base editing was reported in ^1^. The gRNAs used for CRISPRi and CRISPRa of L1FLI-15q21.3 and L1FLI-10q25.1b are listed in Supplementary Table7.

The ABE8.20m, dCas9-VPR and dCas9-KRAB plasmids were linearized with SpeI and purified by phenol-chloroform, as previously described ^3^. IVT of the ABE8.20m, dCas9-VPR and dCas9-KRAB mRNA was performed as in ^3^ and stored at -80 °C in aliquots.

### Primary cell culture and drug treatment

mPB CD34+ HSPC were cultured in serum-free StemSpan SFEM (StemCell Technologies) supplemented with 100 IU ml^−1^ of penicillin, 100 μg ml^−1^ of streptomycin, 2% glutamine, 300 ng ml^−1^ of human stem cell factor (hSCF), 300 ng ml^−1^ of human Fms-related tyrosine kinase 3 ligand (hFLT3L), 100 ng ml^−1^ of human thrombopoietin (hTPO), 1 μM SR1 and 35 nM UM171. Cells were seeded in tissue-cultured treated plates at the concentration of 5x10^5^ ml^-1^ and cultured in a 5% CO_2_ humidified incubator at 37 °C.

For drug treatment experiments 10nM of TAK-243 (MCE) or 100nM of Auranofin (Sigma) were added to the culture media immediately following gene editing, and refreshed every 2 days, unless otherwise specified.

### LNPs preparation

mRNA-LNPs were prepared by rapid pipette mixing. LNPs were formulated by combining an ethanol phase containing the lipid components (ionizable lipid, DOPE, cholesterol, and DMG-PEG2000) at a molar ratio of 46.5:16:35:2.5 with an aqueous phase containing mRNA at an aqueous-to-organic volume ratio of 3:1. The ionizable lipid-to-mRNA weight ratio was maintained at 10:1.

### Gene editing of human HSPC

Between 2.0 × 10^5^ and 7 × 10^5^ mPB HSPC were washed with Ca^2+^and Mg^2+^-free PBS, resuspended in P3 Primary Solution from the P3 Primary Cell 4D-Nucleofector X Kit (Lonza), and electroporated using the program EO-100. Electroporation was performed after 2 or 3 days of culture, as described in the text. For UBA1 editing, cells were electroporated with 100pmol of UBA1 gRNA and 5 μg of ABE8.20-m mRNA. For CRISPRi or CRISPRa of *AAVS1*, 100pmol of AAVS1 gRNA and 5 μg of dCas9-KRAB or dCas9-VPR, respectively, were used. For CRISPRi or CRISPRa of L1FLI-15q21.3 and L1FLI-10q25.1b 50pmol of gRNA 1 and 50pmol of gRNA 2 were electroporated together with 5 μg of dCas9-KRAB or dCas9-VPR, respectively.

For LNPs-mediated gene editing, HSPC were cultured at the concentration of 5x10^5^ ml^-1^ and transduced 3 days after culture (equivalent to 1 day after *UBA1* base editing) with LNPs containing 5ug of dCas9-KRAB or dCas9-VPR mRNA together with 100pmol of AAVS1 gRNA or 50pmol gRNA 1 + 50pmol gRNA2 for L1FLI-15q21.3 and L1FLI-10q25.1b.

After 48h and 96h from electroporation HSPC were collected for RNA analysis, while 7 days after electroporation cells were pelleted for gDNA molecular analysis and evaluation of editing efficiency using Sanger sequencing.

### Clonogenic assay

24h after editing, 500 HSPC ml^-1^ were plated in methylcellulose-based medium (MethoCult H4434, STEMCELL Technologies) supplemented with 100 IU ml−1 of penicillin,100 µg ml−1 of streptomycin, IL-6 (10ng ml−1) and FLT3L (10ng ml−1). Technical duplicates were performed. Colonies were assessed morphologically 14 days after seeding. For experiments using small molecule inhibitors, drugs were added to the methylcellulose-based medium at the following concentrations: TAK-243 10nM, Auranofin 100nM.

### Mice

Animal experiments were approved by the UHN Animal care committee (ARC1117) and performed in accordance with institutional guidelines. 8-12-week-old female NOD.Cg-*Prkdc^scid^Il2rg^tm1Wjl^*Tg(CMV-IL3, CSF2, KITLG)1Eav/MloySzJ (NSG-SGM3) mice were irradiated with 225cGy of Gamma radiation (GammaCell40) and transplanted 24h after with edited HSPC by intrafemoral injection. Mice were euthanized 8 weeks post engraftment by cervical dislocation. Long bones were processed as described in ^4^ and assessed by flow cytometry using the following antibodies: CD14 PE-Cy5 (1:200), GlyA PE (1:100), CD33 BV786 (1:100), CD45 AF700 (1:100), CD34 APC-Cy7 (1:100), CD19 PE (1:100), CD45 V500 (1:50), CD56 BV605 (1:100) and CD3 FITC (1:100). Sytox Blue (1:2000) was used for live and dead staining. Samples were acquired using a BD Symphony A5 (BD Biosciences) flow cytometer calibrated using unstained and single stained controls. Data were analyzed with FlowJo software v.10.5.3 (BD Biosciences).

### *In vitro* multilineage differentiation assay

For in vitro differentiation assay, 1x10^4^ HSPC were seeded 24h after editing in a nontissue culture-treated, 96-well, U-bottomed plate (Falcon) and cultured in StemPro (STEMCELL Technologies) supplemented with StemPro nutrients (STEMCELL Technologies), 100 IU ml^−1^ of penicillin, 100 μg ml^−1^ of streptomycin (GIBCO), 1% glutamine (GIBCO), human LDL (STEMCELL Technologies 50ng ml^-1^), SCF (100 ng ml^-1^), Flt3L (20 ng ml^-1^), TPO (100 ng ml^-1^), IL-6 (50 ng ml^-1^), IL-3 (10 ng ml^-1^), GM-CSF (20 ng ml^-1^) (R&D) and EPO (Jansen, 3 units ml^-1^). The cell culture media was refreshed every 4 days. After 14 days of culture, cells were assessed by flow cytometry using the following antibodies: CD14 BV605, GlyA PE, CD15 PE-Cy5, CD34 APC-Cy7, CD66b AF647, CD45 AF700, CD33 BV786 all at a 1:100 dilutions except CD45RA FITC (1:200). Propidium Iodine (1:5000) was used for live/dead staining. Samples were acquired using a BD Symphony A5 (BD Biosciences) flow cytometer calibrated using unstained and single stained controls. Data were analyzed with FlowJo software v.10.5.3 (BD Biosciences). For experiments using small molecule inhibitors, drugs were added to the culture medium at the following concentrations: TAK-243 10nM, Auranofin 100nM, and refreshed every 2 days.

### Molecular analyses

gDNA was extracted using the QIAamp DNA Micro Kit (QUIAGEN) following manufacturer’s instructions. 50-100ng gDNA were used for PCR amplification of the UBA1 target locus as previously reported ^5^, and the PCR product was purified with the QUIaquick PCR Purification Kit (QUIAGEN) according to manufacturer’s instructions. Editing efficiency at the UBA1 locus was quantified by Sanger sequencing and analyzed using EditR software (http://baseeditr.com) with default parameters.

RNA was extracted using the RNeasy Micro Kit (QUIAGEN) and RNase-free DNAse set (QUIAGEN) following manufacturer’s instructions. RNA was reverse transcribed into cDNA according to the SuperScript IV VILO with ExDNAse Kit (Thermo Fisher Scientific) protocol. 3ng of cDNA were used for gene expression analysis by ddPCR (QX200, Biorad) following the thermal cycling protocol recommended by manufacturer’s instructions and results were analyzed using QuantaSoft Software (v1.7.4 Biorad). Target gene expression of SPI1 (Hs02786711_m1, ThermoFisher) and IRF1 (Hs00971962_g1, ThermoFisher) was determined normalizing on HPRT1 expression (Hs02800695_m1, ThermoFisher) and represented as a fold change relative to the control condition.

### scRNA-seq analysis of public datasets and TE quantification

The combined scRNA-seq of BM CD34+ HSPC form Molteni et al. ^5^ (GEO accession GSE272816), elderly samples from Ainciburu et al. ^6^ (GEO accession GSE180298), and CD34+ sorted samples from Wu et al. ^7^ (GEO accession GSE190652,) was integrated and processed as described in ^5^. The combined Healthy vs Young scRNAseq and the MDS vs Aged Healthy HSC were obtained from ^6^, and the combined HSC-iM/HSC-I scRNAseq was obtained from ^8^. To maintain consistency with the VEXAS scRNAseq, cells with <200 genes or >8000 genes or >20% mitochondrial genes were excluded to remove dying cells and doublets. scRNAseq processing was performed in R using Seurat. Samples were LogNormalized and scaled by a factor of 10.000, and the top 20% most variable genes were selected for analysis. Cell type annotation and pseudotime analysis were performed using the Bone Marrow reference map published in ^9^ (https://github.com/andygxzeng/BoneMarrowMap).

Transposable elements contained in the highly curated Telescope annotation file belonging to HERVs and L1 families were profiled at locus resolution (hg38) using Stellarscope ^10^ v 1.5) with default parameters, restricting the quantification to cells passing QC from the scRNA-seq analysis and using pooling_mode “individual” and reassigning_mode “best_exclude". To avoid having the same UMI contributing to both gene expression counts and transposable element counts matrices, stellascope_resolve was used to obtain a TE exclusive count matrix, which was used for subsequent analysis. Data integration from TE counts was performed in R (4.3.1) using Seurat (v. 5.2.1), LogNormalized and scaled to a factor of 10.000. Batch correction was performed for donor and dataset effects with the Harmony package as described in ^5^.

Co-expression gene networks in VEXAS patients HSC and healthy donor HSC were obtained through hdWGCNA ^11^, with default parameters. Gene hubs were calculated for each program considering the top 50 interconnected genes. Transcriptional program conservation across datasets was performed following the WGCNA module conservation analysis, with default parameters. Pathway enrichment analysis was calculated for each hdWGCNA transcriptional program using gProfiler ^12^ with default parameters.

### Differential TE expression

Differentially expressed TE between VEXAS patients and controls in each cell type were obtained using Find Markers from Seurat (test “MAST”) with the following parameters: min.pct = 0.1, only.pos = FALSE, test.use = MAST, logfc.threshold = 0.25. TE with padj < 0.05 were considered differentially expressed. Cell types were defined as in ^5^ and only populations with at least 50 cells in both VEXAS and HD conditions were considered.The VEXAS TE primitive score was calculated as pseudotime-weighted average difference between the log2FC expression of each of the 21 TE differentially expressed in VEXAS HSC vs controls in HSC minus the log2FC expression of the same TE in each downstream HSPC population. The VEXAS TE unique score was calculated as average difference between the log2FC expression of each of the 21 TE differentially expressed in VEXAS HSC vs control HSC and the log2FC of the same TE in MDS HSC vs control HSC from ^6^, aged HSC vs young HSC from ^6^ or HSC-iM vs HSC-I from ^8^.

TE-WGCNA correlation analysis was computed as reported in ^13^. Briefly, expression correlation was quantified using Spearman and results were considered significant if |rho| > 0.2, padj < 0.05 and padj bootstrap_1000_ < 0.05. For TE-TF correlation analysis, only TF with at least one motif binding site (predicted by FIMO) within the TE loci alongside a TE-TF Spearman |rho| > 0.2, padj < 0.05 and padj bootstrap_1000_ < 0.05 were considered significant. Customized networks were generated using the network R package (v.1.19.0). Customized dotplots and heatmaps were generated using the ggplot2 R package (v.4.0.2).

### Quantifications and statistical analyses

Biological replicates are indicated by ‘n’. For the in vivo experiments, multiple mPB donors were pooled to reduce donor-driven variability and to increase cell numbers. Unless otherwise specified, data are presented as the median with the IQR (or range) or the mean ± s.e.m. depending on data distribution. Unless otherwise noted, A Geisser-Greenhouse corrected One Way Anova with multiple comparisons was performed for *in vitro* experiments. Statistical analysis of *in vivo* experiments was performed using the Mann-Whitney U-test between each pairwise comparison. For all analyses the significance threshold was set at 0.05: \**P* < 0.05; \*\**P* < 0.01, \*\*\**P* < 0.001, \*\*\*\**P* < 0.0001. GraphPad Prism v.10.2.3 was used for statistical analysis. Only the most relevant pairwise comparisons are shown in the figures. Unless otherwise noted, non significant comparisons were not shown.

## Acknowledgments

We thank patients and their families for their consent to donate samples for research; the Leukemia Tissue Bank at Princess Margaret Cancer Centre for sample coordination; Attya Omer for the dCas9-VPR and dCas9-KRAB construct design and sharing; Mark D. Minden and Andrea Arruda for providing mPB samples.

## Author contributions

A.V. performed research and interpreted data. S.D. performed experiments under the supervision of B.L. C.G., K.K., T.V., and L.J. performed research. J.McL. and I.D.B. performed research under the supervision of S.Z.X. I.M., I.N.-B., and J.M. contributed new reagents/analytic tools. A.G.X.Z. provided bioinformatic support. J.E.D. and S.F. supervised the research. J.E.D. obtained funding. A.V. prepared and drafted the manuscript and S.Z.X, K.K., S.F. and J.E.D. contributed to manuscript organization and editing. All authors proofread the manuscript.

## Data availability

All relevant data are included in the paper and/or its supplementary information files. The scRNAseq datasets used in these studies were downloaded from GEO (GSE272816, GSE196052, GSE180298, GSE249479, GSE196990) and EGA (EGAS00001007358).

## Code availability

Custom code underlying key analyses in the main figures are available upon request.

## Corresponding author

John E. Dick

**Correspondence and requests for materials** should be addressed to J.E.D.

## Supplementary Information

Supplementary Tables 1-7

Extended Data Figures 1-6

## Extended Data Figure Legends

**Extended Data Fig.1.**
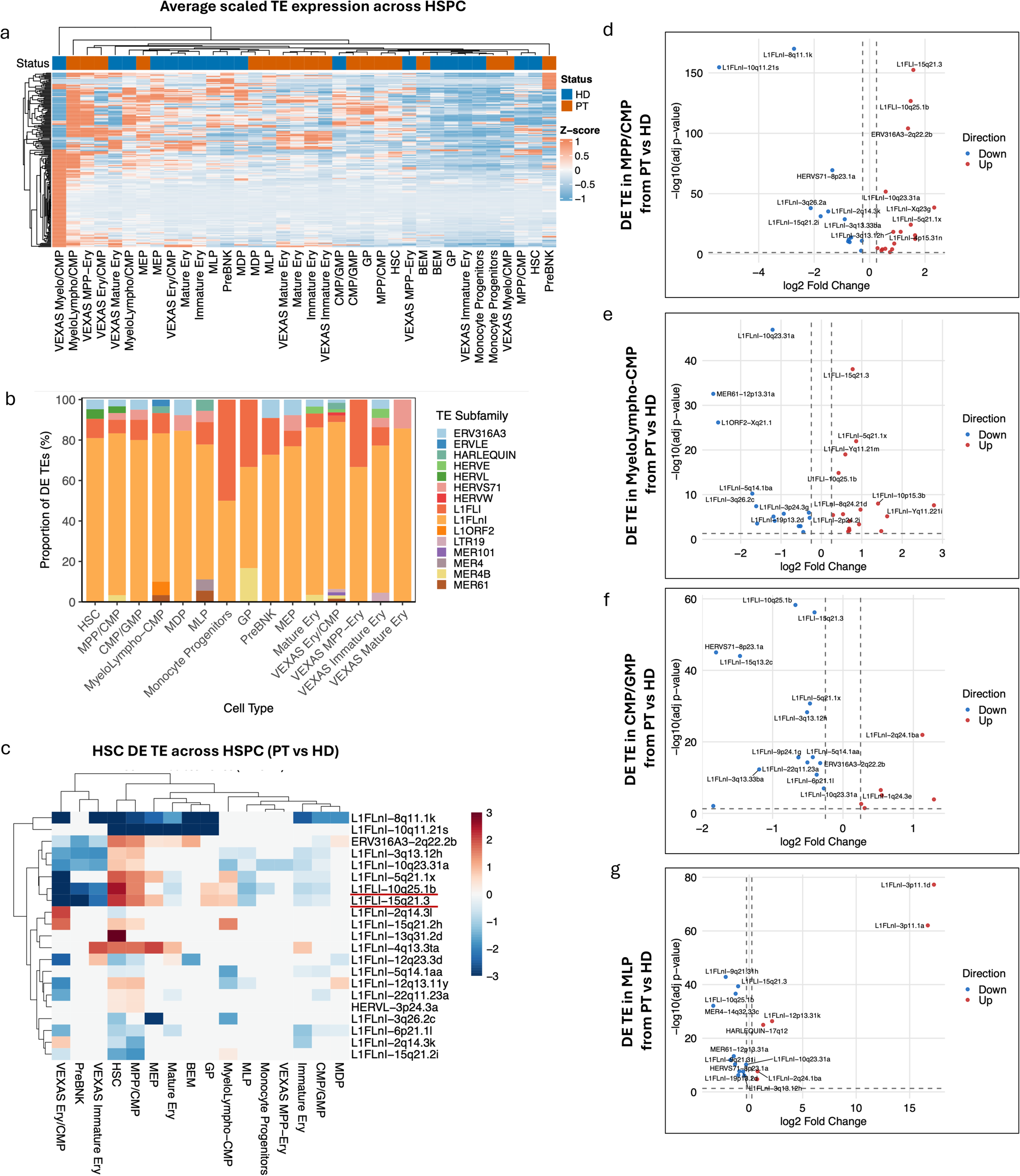
Differential TE expression across VEXAS HSPC relative to HD. a) Heatmap showing the average scaled TE expression of the top 200 most variable features across VEXAS and HD HSPC cell types defined in <u>(Molteni et al. 2025)</u>. b) Bar plot showing the proportion of differentially expressed TE loci by TE family across HSPC populations in VEXAS patients versus HD. c) Heatmap showing the average log2 fold-change of the 21 TE loci identified in Fig. 1c across HSPC populations in VEXAS patients relative to HD. d-g) Volcano plots showing upregulated (red) and downregulated (blue) TE loci in (d) MPP/CMP, (e) Myelo-Lympho-CMP, (f) CMP/GMP and (g) MLP from VEXAS patients versus HD.

**Extended Data Fig.2.**
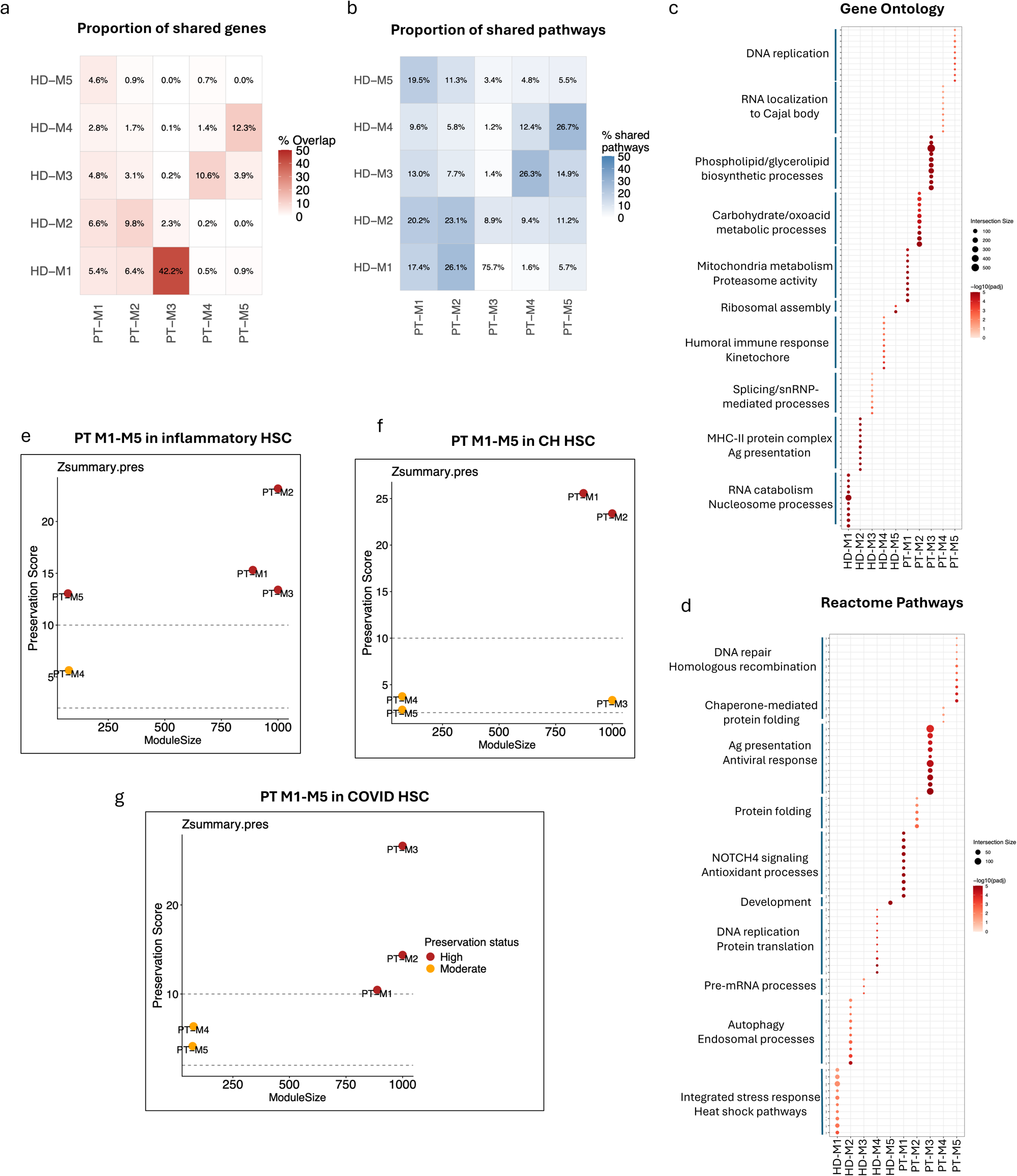
VEXAS HSC transcriptional programs are shared with HSC from related hematological conditions. a) Heatmap showing the proportion of shared genes between hdWGCNA modules from HSC of HD (HD-M1 to HD-M5) and VEXAS patients (PT-M1 to PT-M5). b) Heatmap showing the proportion of shared pathways identified by g:Profiler enrichment analysis of hdWGCNA gene modules in HSC from HD (HD-M1 to HD-M5) and VEXAS patients (PT-M1 to PT-M5)., d) Unique (c) Gene ontology and (d) Reactome terms enriched by g:Profiler enrichment analysis of hdWGCNA gene modules in HSC from HD (HD-M1 to HD-M5) and VEXAS patients (PT-M1 to PT-M5). Up to 10 pathways per each module are shown. e-g) hdWGCNA module preservation analysis comparing VEXAS HSC gene modules in (e) inflammatory HSC ^3^, (f) HSC from clonal hematopoiesis donors ^2^ and (g) HSC form COVID-19 patients ^31^.

**Extended Data Fig.3.**
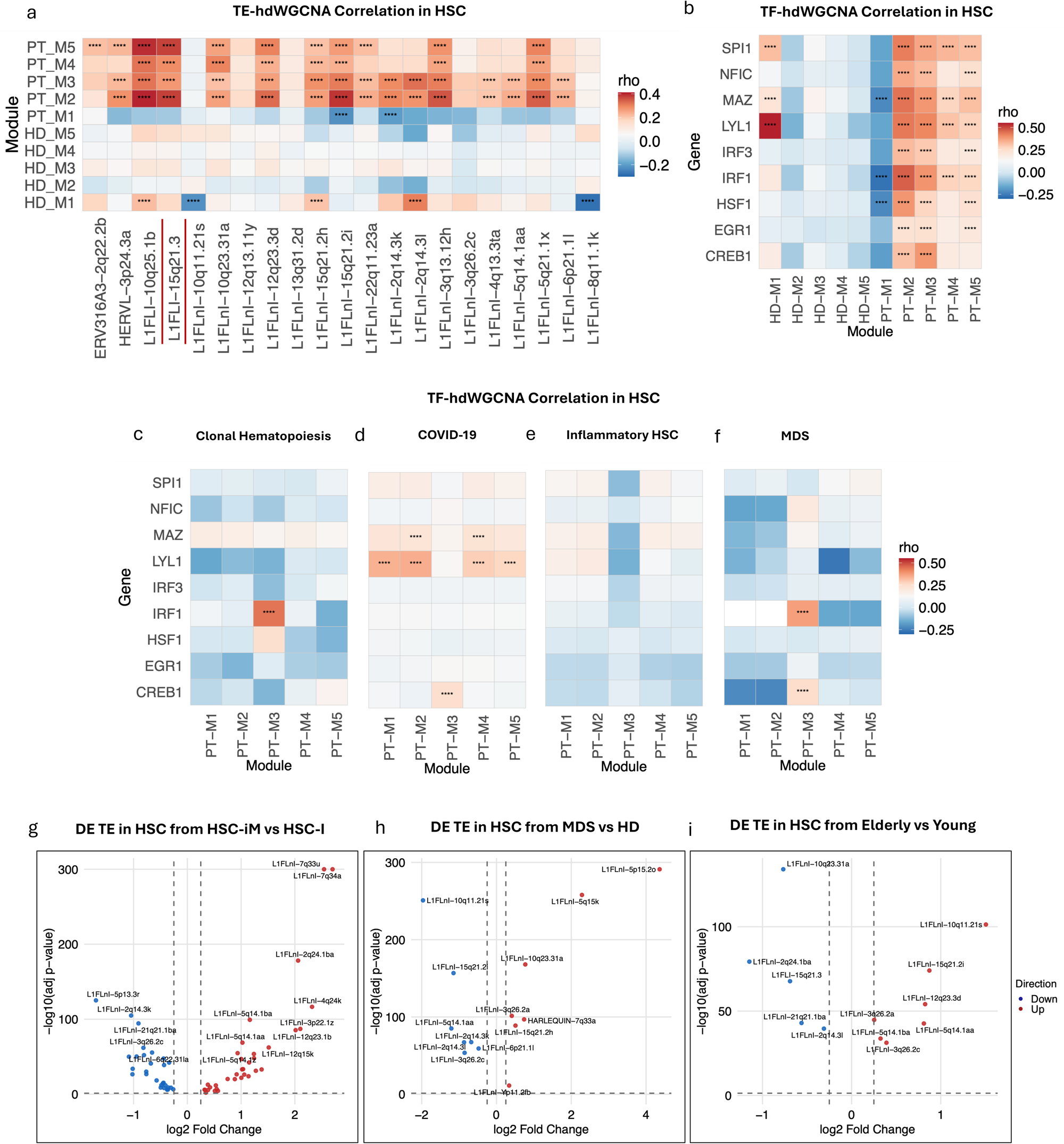
Correlative expression between TE - TF associations and transcriptional programs in HSC from various hematological conditions. a,b) Heatmap showing Spearman correlation between (a) TE expression or (b) TF expression and hdWGCNA gene modules in HSC from VEXAS patients and HD.c-f) Heatmap showing Spearman correlation between TF expression and hdWGCNA gene modules in HSC from (c) clonal hematopoiesis donors ^2^, (d) COVID-19 patients ^31^, (e) xenografts subjected to acute inflammatory challenge ^3^, and (f) MDS patients ^28^. g-i) Volcano plots showing upregulated (red) and downregulated (blue) TE loci in (g) HSC-iM relative to HSC-I, (h) MDS HSC relative to healthy donor HSC and (i) Elderly HSC relative to young HSC.

**Extended Data Fig.4.**
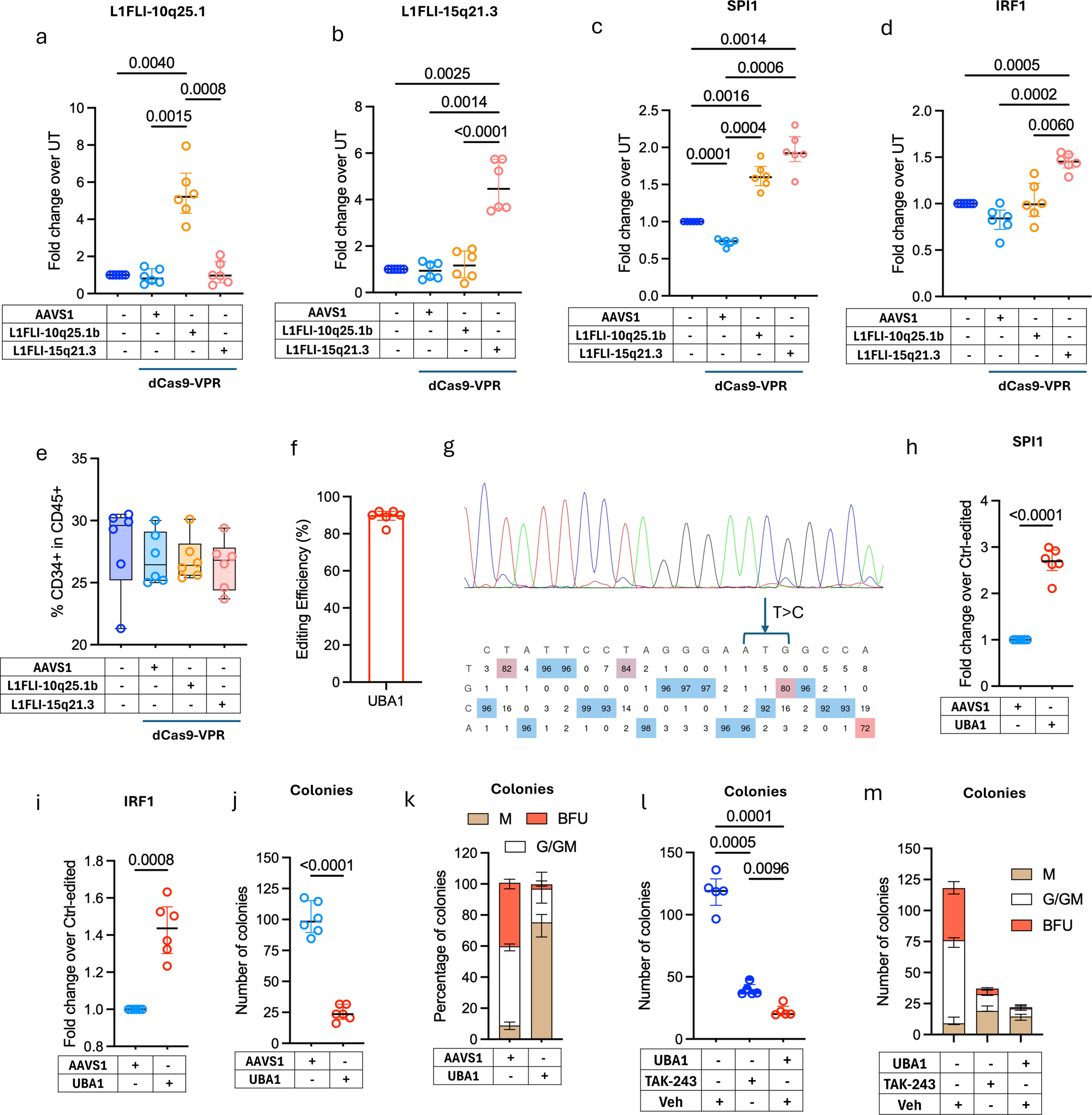
L1-10 and L1-15 transactivation phenocopies *UBA1* mutational HSPC phenotypes. a-d) Fold change in expression of (a) L1-10, (b) L1-15, (c) SPI1, and (d) IRF1 of HSPC edited as indicated relative to UT 48 h after editing, measured by ddPCR (n=6). e) Proportion of CD34+ cells within CD45+ cells measured by flow cytometry of HSPC edited as indicated after two weeks of differentiation culture (n=6). f) Percentage of HSPC carrying the intended Met41Thr mutation at the *UBA1* gene 7 days after editing (n=6). g) Representative plot of *UBA1* Sanger sequencing. h,i)) Fold change in expression of (h) SPI1 and (i) IRF of *UBA1* mutant HSPC relative to control edited cells 48 h after editing, measured by ddPCR (n=6). j) Number of colonies obtained on seeding HSPC edited as indicated (n=6). k) Proportion of myeloid (M), granulocytic/granulocytic-myeloid (G/GM) and erythroid (BFU) colonies from the condition in (j) (n=6). l) Number of colonies obtained on seeding HSPC edited or treated as indicated (n=6). m) Proportion of myeloid (M), granulocytic/granulocytic-myeloid (G/GM) and erythroid (BFU) colonies from the condition in (l) (n=6).

**Extended Data Fig.5.**
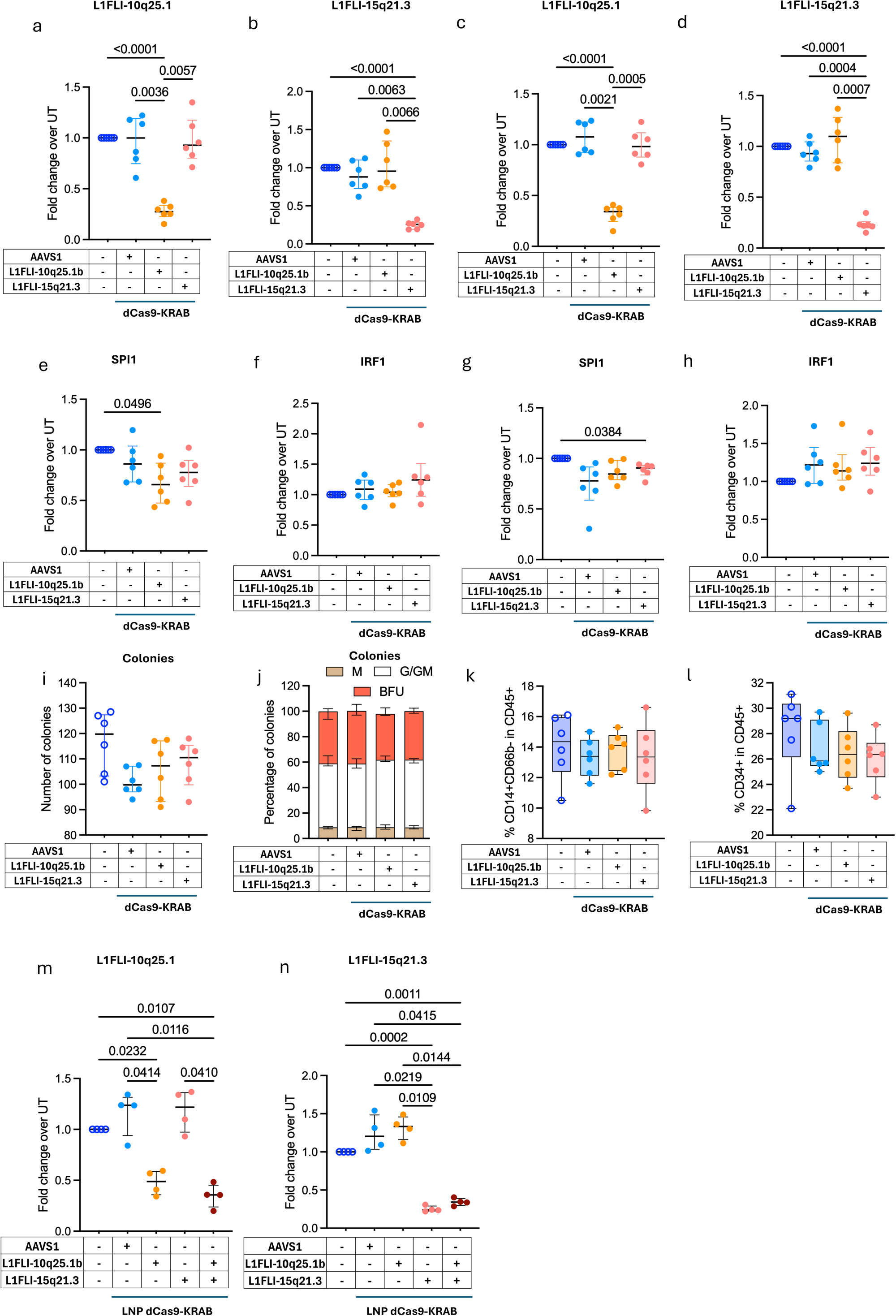
L1-10 and L1-15 silencing does not impact healthy HSPC. a-h) Fold change in expression of (a,c) L1-10, (b,d) L1-15, (e,g) SPI1 and (h,i) IRF1 of HSPC edited as indicated relative to UT (a,b,e,f) 48 h or (c,d,g,h) 96 h after editing measured by ddPCR (n=6). i) Number of colonies obtained on seeding HSPC edited as indicated (n=6). j) Proportion of myeloid (M), granulocytic/granulocytic-myeloid (G/GM) and erythroid (BFU) colonies from the condition in (i) (n=6). g,h) Proportion of (g) CD14+CD66b- and (h) CD34+ cells within CD45+ cells measured by flow cytometry of HSPC edited as indicated after two weeks of differentiation culture (n=6). m,n) Fold change in expression of (m) L1-10 and (n) L1-15 of HSPC edited as indicated relative to untreated cells 96 h after editing measured by ddPCR (n=4).

**Extended Data Fig.6.**
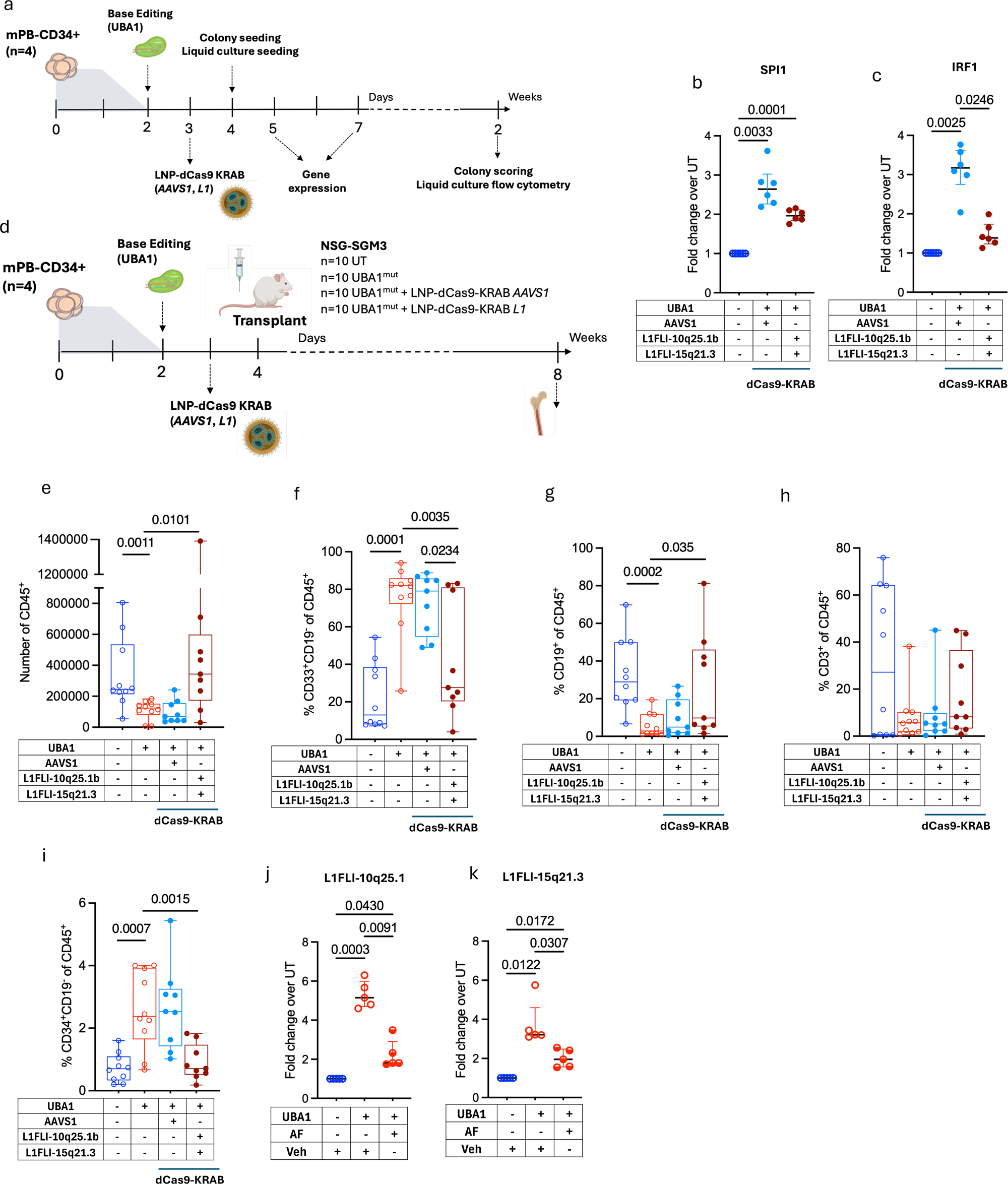
L1-10 and L1-15 silencing in *UBA1* mutant HSPC mitigates VEXAS phenotypes. a) Scheme of the double gene engineering editing protocol. b,c) Fold change in expression of (b) SPI1 and (c) IRF1 of HSPC edited as indicated relative to untreated cells 96 h after editing measured by ddPCR (n=6). d) Scheme of *in vivo* experiments. e) Number of CD45+ human cells per femur of xenografted mice at 8w postengraftment. f-i) Percentage of (f) CD33+CD19- (g) CD19+ (h) CD3+ and (i) CD34+CD19-cells within CD45+ human cells of xenografted mice at 8w postengraftment. j,k) Fold change in expression of (j) L1-10 and (k) L1-15 of *UBA1* mutant HSPC and *UBA1* mutant HSPC treated with Auranofin (AF, 100nM) relative to vehicle treated cells (n=5).

